# Metabolomic and transcriptomic analyses reveal the effects of grafting on anthocyanin synthesis in grapevine

**DOI:** 10.1101/2021.10.09.463741

**Authors:** Haixia Zhong, Zhongjie Liu, Fuchun Zhang, Xiaoming Zhou, Xiaoxia Sun, Wenwen Liu, Hua Xiao, Nan Wang, Mingqi Pan, Xinyu Wu, Yongfeng Zhou

**Author notes:** These authors contributed equally to this work (Haixia Zhong, Zhongjie Liu, Fuchun Zhang). Corresponding authors: Mingqi Pan,; Xinyu Wu,; Yongfeng Zhou.

## Abstract

The grafting has been commonly used in viticulture, which joints the scion from a cultivar with the stem of a rootstock. Grafting has crucial impacts on various phenotypes of the cultivar including berry metabolome and berry coloring, however, the genetics and regulation mechanisms are largely unexplored. In this study, we analyzed the phenotypic, metabolomic and transcriptomic profiles at three stages (45, 75 and105 days after flowering) of the Crimson Seedless (Vitis vinifera, CS) cultivar grafted to four rootstocks (three heterografting: CS/101-14MG, CS/SO4, CS/110R and one self-grafting CS/CS) with an own-rooted grafting-free Crimson Seedless (CS) as a control. All the heterografting plants had a significant influence on berry reddening as early as ~45 days after flowering. The grafting of rootstocks promoted anthocyanin synthesis and accumulation in grape berries. The metabolomic features showed that Cyanidin 3-O-glucoside, Delphinidin 3-O-glucosid, Malvidin 3-O-glucoside, Peonidin 3-O-glucoside and Petunidin 3-O-glucoside were the pigments responsible for the purplish-red color peels. Transcriptomic analyses revealed that the anthocyanins biosynthetic related genes from the upstream (phenylalanine ammonia-lyase) to the downstream (anthocyanidin 3-O-glucosyltransferase and anthocyanidin synthase) were upregulated with the accumulations of anthocyanins in CS/101-14MG, CS/SO4 and CS/110R. At the same time, all these genes were also highly expressed and more anthocyanin was accumulated in CS/CS samples compared to CS samples, suggesting that self-grafting rootstocks might also have promoted berry reddening in grapevine. Our results provide global transcriptomic and metabolomic features in berry coloring regulation under different grafting conditions for improving the berry quality in grapevine production.

## Introduction

The grafting had been practiced in horticultural plants ~4000 years ago in China^1^, which established a vascular continuity by joining the scion of one plant with the stock of another plant. The rootstocks could benefit the scion plant on enhancing the resistance to biotic and abiotic stresses, and elevating desired agronomic traits. The stem grafting in grapevine production could be traced back to ~2500 years ago^2^. The practice of grafting in grapevine using the wild vitis species as rootstocks, which bring advantages to the scion cultivar, including flowering time, berry quality, dwarfing, disease or pest resistance and environmental adaptation.

Grapevine coloring is a very important agronomic trait that are required for adaptation to the markets including table grapes and wine making. There are two types of grape berry coloring: peel coloring and flesh coloring. In general, most red grapes are pigmented in the peel, and the accumulation of anthocyanins in ripening grape berries only occurs in epidermal and subepidermal cells^4,5^. Recent studies revealed that the grape peel color was mainly determined by the composition and content of anthocyanins^6^, and the relative proportion of anthocyanins in each grape variety is stable^7^. The anthocyanins in grapes mainly include anthocyanin, delphinidin, petunidin, peonidin and malvidin, which are composed of aminoglycosides or glycosides and acylation^16^. In grapevine, the content of anthocyanins in interspecific hybrids were lower than the wild *Vitis* species, and table grapes were lower than wine grapes^17^. Moreover, the biosynthesis of anthocyanins is affected by light^8^, temperature^9^, moisture^10^, mineral nutrients^11^, cultivation measures^12,13^, growth regulators^14,15^ and other external factors. The VvMYBA1 binds to VvWDR1 and activates three promoters *(VvCHI3, VvOMT,* and *VvGST4),* which positively regulates berry flesh color, while *VvMYBC2-L1* negatively regulates this process by competing the binding site with the R2R3-MYB transcriptional activators or by repressing the expression level of *VvOMT* and *VvGST4*^17^ Genomic structural variants showed that the QTL region underlying berry color is hemizygous and convergent evolution was associated with the origin of the green coloring in grapevine^18^. It is known that grafting connects two different genomes and introduces complex genomic regulations in long-living perennials^3^. A subclade of *β*-1,4-glucanases contributed to the grafting among a tomato scion, a *Nicotiana benthamiana* middleman and an *Arabidopsis* rootstock by facilitating cell wall reconstruction^19^. In addition, heterografting by using the scion of sweet orange and rootstock *P. trifoliata* was performed to investigate the sRNA-mediated graft-transmissible epigenetic modifications in citrus grafting^20^. Rootstock influenced the pigment on grape peel of scion cultivar was overserved^21^. However, the genetic basis and molecular mechanism of effects of grafting on grape peel color is still unknown.

Crimson seedless grape is an important grape cultivar with bright red fruit grains, and yellow flesh. It’s natural seedless late-ripening European subspecies with thick fruit powder, translucent flesh, hard flesh, high content of soluble solids. Understanding the effects of grafting on anthocyanin synthesis pathway could be valuable for grapevine production.

In our study, we aimed at understanding metabolic differences and significantly differentially expressed genes in anthocyanin biosynthesis during berry development in heterografting (CS/101-14MG, CS/SO4, CS/110R), self-grafting (CS/CS) and grafting-free (CS) plants. We studied the association of grafting, berry coloring, metabolomic and transcriptomic profiles and found the hub genes play critical roles in anthocyanin biosynthesis.

## Materials and methods

### Plant materials and treatments

The grafting experiment was performed at the Xinjiang academy of agricultural sciences anningqu comprehensive test field, national grape industry technology system fruit quality control post base, Xinjiang, China. Scions were selected from thrive annual branches on Crimson seedless self-root grapevine (CS). Four grafting combinations were constructed: one with 101-14MG, (CS/101-14MG) grafted as rootstock, one with SO4 (CS/SO4) grafted as rootstock, one with 110R (CS/110R) grafted as rootstock and one with Crimson seedless (CS/CS) grafted as rootstock (Figure. 1). Every grafting combination was performed more than ten repeats. Berry skins were collected at three stages: 45, 75 and105 days after flowering (DAF) (Figure. 1). All samples with at least 50 berries were collected in randomized block designs and three biological repeat. After being taken back to the laboratory and the peels were carefully excised, and then collected and frozen in liquid nitrogen. After being roughly ground, a total of 45 samples were stored at −80 °C for metabolome, mRNA sequencing and RT-qPCR validation.

**Figure 1:**
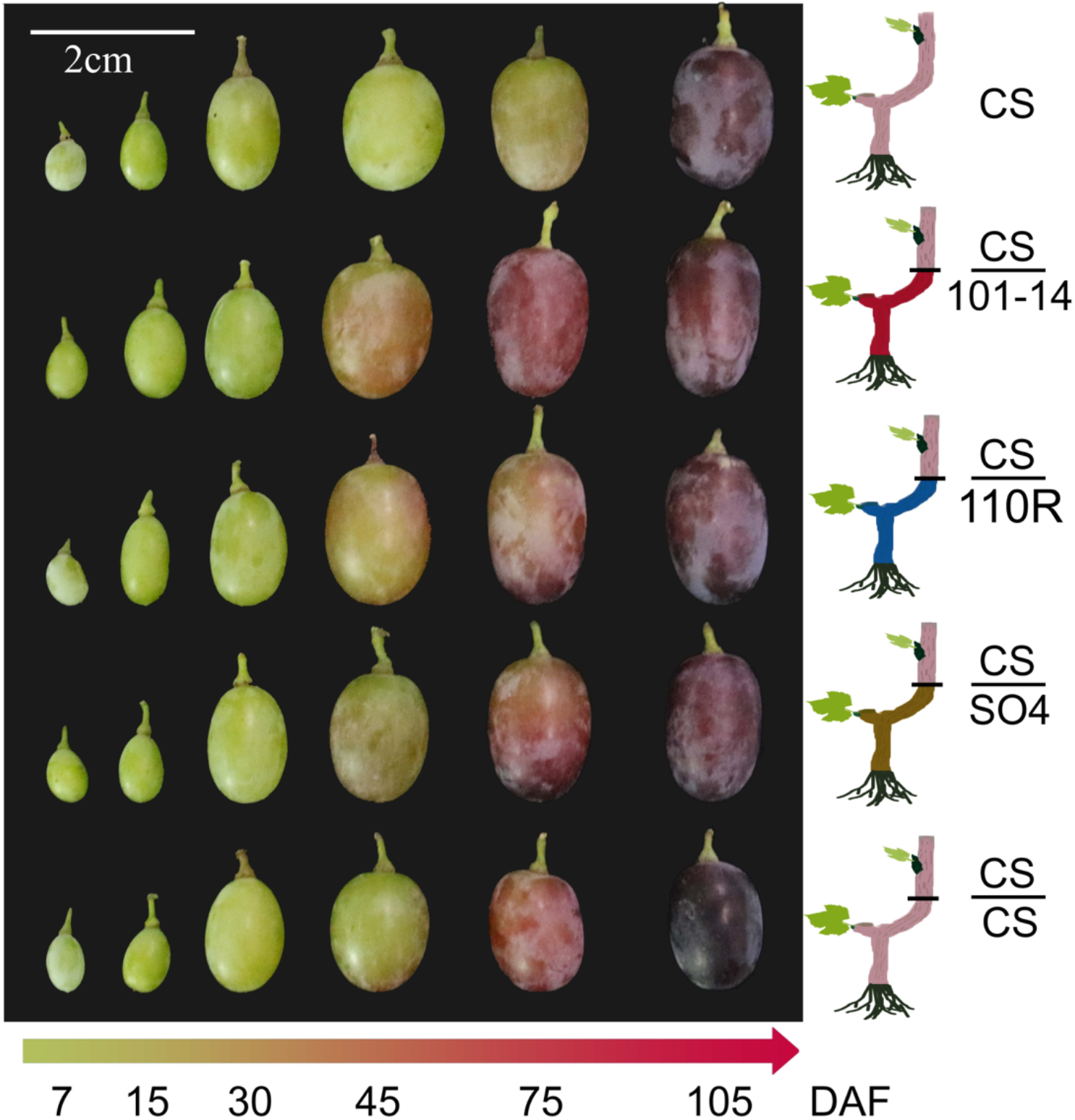
The grafting design and berry phenotypes of Crimson Seedless grafted on different rootstocks. A schematic illustration of the grafting and Phenotypes of grape berry in 6 development stages.

### Metabolite identification and quantification

The anthocyanins profiles for each sample were conducted in the following three steps: grinding, extraction and measurement. (i) Using mixer mill (MM 400, Retsch) to crush the freeze-dried sample. (ii) The 50mg powder with extracting solution (methanol: water: hydrochloric acid, 799:200:1, V/V/V) vortexed and ultrasound 10 min separately, then centrifuged at 12, 000 g and 4 °C for 3 min, and collect the supernatants. The precipitate was treated again using the same method to fully extract the components. Combine the supernatants and filtrated (PTFE, 0.22 μm; Anpel) for UPLC-MS/MS analysis. (iii) UPLC (ExionLC™ AD) and Tandem Mass Spectrometry (MS/MS) (QTRAP^®^ 6500+, N) used to detected the contents of Anthocyanins. Substituting the integral peak area of all the detected samples into the linear equation of the standard curve for calculation, and further putting it into the calculation formula, the absolute content data of the substance in the actual sample is finally obtained.

### RNA-Seq and analysis of differentially expressed genes (DEGs)

The total RNA was isolated by proceed as following: (i) add the preheated cracking liquid and β-Mercaptoethanol; (ii) add equal volume chloroform / isoamyl alcohol (24 / 1); (iii) shaking and centrifugation, take the supernatant, add equal volume chloroform / isoamyl alcohol (24/1) and then centrifugation; (iv) repeat iii again, add precipitant for precipitation and centrifugation and wash with ethanol and recover RNA. The obtained RNA was handed over to Shanghai Personal Biotechnology Cp. Ltd for making library and RNA-sequencing.

By using fastp^29^ with default parameters, the high-quality clean reads were filtered from the raw reads). Then, the clean reads were aligned to the *Vitis vinifera* reference genome (12X, http://plants.ensembl.org/Vitis_vinifera/Info/Index) using HISAT2^30^. The mapped reads were assembled using the software StringTie^31^(http://ccb.jhu.edu/software/stringtie/), then the read count value of each mapped gene counted by using HTSeq^32^ as the original expression level of the gene, and FPKM was used to normalized the expression level. Genes with | log2FoldChange | > 1 and significant P-value < 0.05 calculated by DESeq^33^ were identified as differentially expressed.

The principal component analysis (PCA) was used to find associations in the metabolome and transcriptome data set and revealed specific metabolite and transcripts in categories^34–36^. The results were analyzed and visualized using R Studio software (https://www.rstudio.com/) and two packages FactoMineR and factoextra.

### The enrichment analysis of gene function

We used ClueGo+Cluepedia in Cytoscape^37^ to classify genes functionally, and merge related terms that share similar related genes to reduce redundancy. The GO-term fusion function with default parameters was used to fuse similar items, and the threshold P<0.05. Use Benjamini and Hochberg’s FDR for hypergeometric testing. Kappa scores were used to group terms using default parameters. The Cytoscape and R were used to visualize the results.

### **The hub gene identification using** the WGCNA analyses

The Weighted Correlation Network Analysis (WGCNA)^38^was used for detecting the hub genes. Firstly, the cluster analysis was performed on the samples according to the expression levels of all genes, Then the TomSimilarity module was used to calculate the co-expression similarity coefficient among genes. To realize the functional connection of genes, the PickSoftThreshold function of the software package was used to select the parameters and carry out the weighted calculation to convert the expression similarity coefficient of the intermediate parameters into the connection between genes. The POWER value was selected when the correlation coefficient tends to be stable. According to the network construction parameters selected above, a weighted co-expression network model was established to classify genes and divide thousands of genes into several modules. After the module is obtained, the gene expression in the module is used to calculate the characteristic gene (ME) of the module, or the first main component of the module. The correlation between the characteristic gene of the module and the trait was further calculated, including the correlation between the gene and the characteristic expression in the module (module membership, MM), the correlation between each gene and the target trait (gene significance, GS).

Following a previous study^38^, we used a passing threshold: GS.abs > 0.5 and GS.pvalue < 0.001 to get genes or modules with significant correlation with traits, and a passing threshold: GS.abs > 0.5 and MM .abs > 0.8 to get the hub gene of each module. The transcription factor annotations were searched for the hub genes using PlantTFDB^39^ (v5.0, http://planttfdb.gao-lab.org/). The cytoscape software was used to visualize the gene interaction network.

### RT-qPCR validation

The extraction and quality detection of RNA used for RT-qPCR and RNA-Seq were carried out in the same batch. Primer3 (v4.0, https://bioinfo.ut.ee/primer3-0.4.0/) and NCBI Primer-BLAST (https://www.ncbi.nlm.nih.gov/tools/primer-blast/index.cgi) were used to design primers for RT-qPCR analyses (Table S1). The *VvGADPH* gene was selected as the housekeeping gene to correct and compute the relative expression of other genes. The PCR assay was performed according to the following conditions: (i) 95 °C for 2 minutes; (ii) 40 cycles at 95 *°C* for 5 seconds, 60°C for 30 seconds, and 72 °C for 10 seconds; (iii) 72 °C for 10 min.

## Results

### Berry development and coloring

We collected berry skin samples from Crimson-Seedless self-rooted and grafted on 4 different rootstocks, including three widely used commercial varieties, and one Crimson-Seedless itself to erase the influence of grafting. Berry on grapes grafted with the three commercial rootstocks showed an earlier start (38.5 days after flowering) of berry coloring and bigger fruit size than CS and CS/CS, in which the heterografting plant CS/101-14MG, performed most obvious (Figure 1). Under the constant observation, three developmental periods were identified based on phenotypic features and color changes. The first stage identified was 45 days after flowering (DAF), in which the skin of the three heterografting grafted samples (CS/101-14MG, CS/SO4 and CS/110R) showed visible color but no difference in fruit size (Figure 1). At 75 DAF, All samples are going to veraison with the berry skin turn to red except CS, and the fruit size of the heterografting samples (CS/101-14MG, CS/SO4 and CS/110R) is bigger than self-grafting CS/CS and grafting-free CS. At the final stage (105 DAF), all samples finished veraison, in which the CS/CS showed the darkest red and smallest fruit size, and the fruit size in sample CS/101-14MG, was biggest, which suggested all the three commercial rootstocks promote fruit development and the best rootstock is 101-14 (Figure 1).

### The metabolomic analyses detected metabolites related to anthocyanin synthesis

The metabolome of a total of 45 samples from five groups (at three stages with three replicates each) of grapevine plants were evaluated, and thirty kinds of metabolites related anthocyanins were identified and classified in seven groups, including Cyanidin, Procyanidin, Peonidin, Delphinidin, Malvidin, Pelargonidin and Petunidin (Figure 2). The content of 26 metabolites (86.7%) was increasing during the development process, which showed a strong correlation with the berry coloring phenotypes. The association of phenotype and metabolomics revealed a capture of critical period associated with the grafting (Figure 2). The unsupervised multivariate principal component analysis of the metabolites showed the first three principal components explained 74.4% of the variance, while PC1 (34.4%) and PC2 (27.2%) described the compounds distribution of samples (Figure 2A and 2B). The results revealed that the increased content primarily showed a low level but gradually accumulated until the highest in third stage. There are 18 and five compounds, mostly colored anthocyanins, explained better in PC1 and PC2, respectively (Figure 2A and 2B, Variable correlation > 0.6). Three Procyanidins and two Cyanidins were separated by PC3, showed a decreasing pattern during the development of grafting.

**Figure 2.**
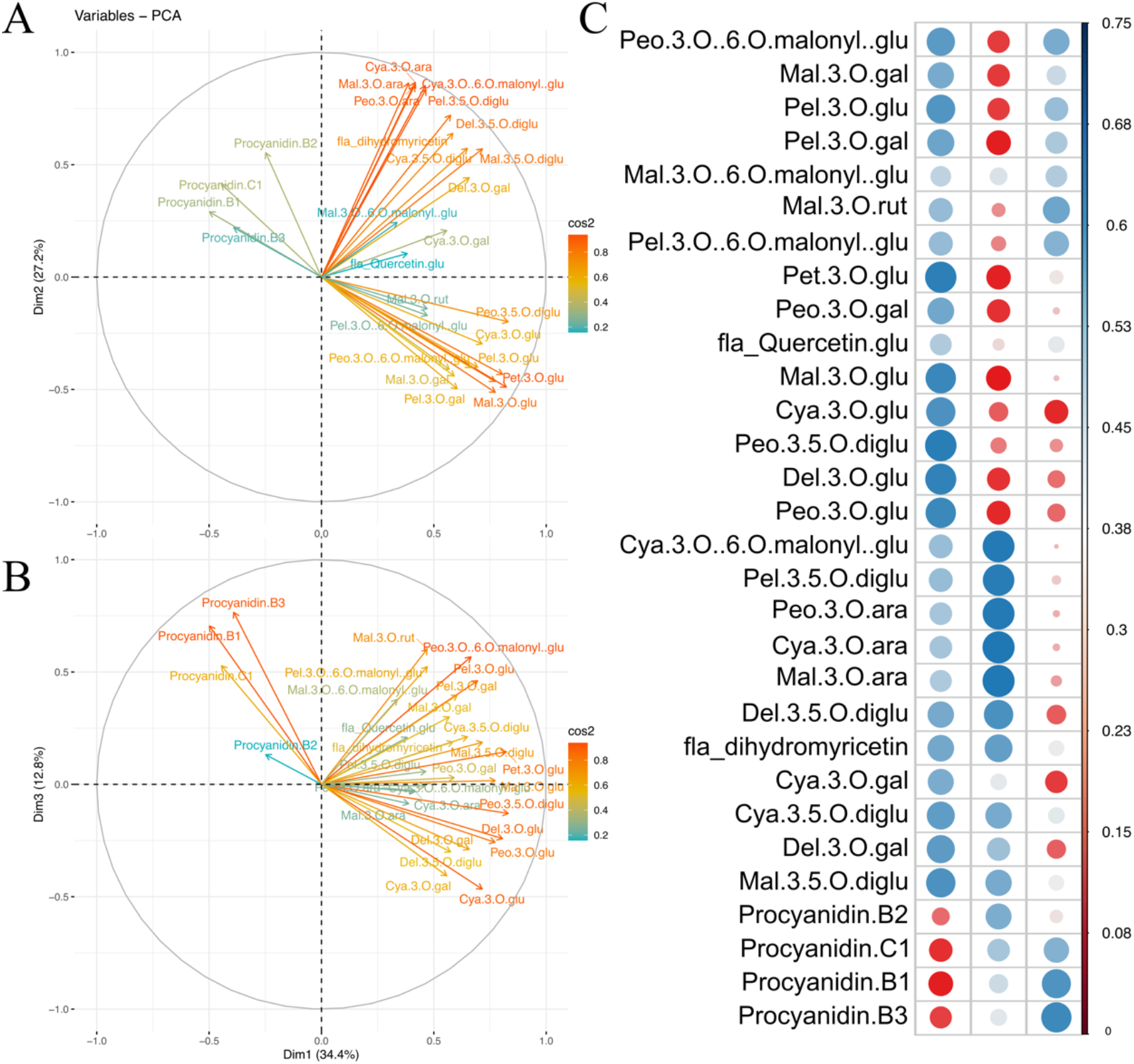
The unsupervised multivariate PCA analyses of the metabolites and its association with berry coloring. The variable correlation plots of 30 metabolites, the distance between variables and the origin measures the quality of the variables on the factor map, and colored by cos2 value (A and B). The Heatmap of cos2 of variables on all the dimensions (C).

### An overview of the transcriptomic data

A total of 1.89 billion clean paired-end reads with a length of 150 bp were obtained from the RNA-sequencing dataset of 45 samples. All clean reads were mapped to the PN40024 reference genome (Ensembl; *Vitis vinifera* 12X). The uniquely mapped rate was > 90% in all samples (Table S2). PCA was used to visualize and evaluate the overall changes in gene expression on different drafting situation. First two PC explained 86.2% of the variation, the first PC (72.6%) separated samples according to development stages for all samples, and the second PC (13.6%) separated the self-grafting CS/CS and the other four groups of samples (Figure 3A). According to the PCA, the distance between the three replicates of each sample was close, suggesting the data is of high quality. In addition, the five samples at 45 DAF showed the similar PC1 value from −25~-50, at 75 DAF PC1 value of the three heterografting samples (CS/101-14MG, CS/SO4, and CS/110R) were around 25, while the PC1 value of self-grafting CS/CS and grafting-free CS only were < −20, and at 105 DAF the three heterografting and grafting-free samples gathered a on the far right while the self-grafting sample CS/CS had the most significant changes compared to 75 DAF. It revealed that the significant transcriptional changes of three heterografting rootstock varieties compared with the two control samples occurred mainly in the second stages (75 DAF).

**Figure 3.**
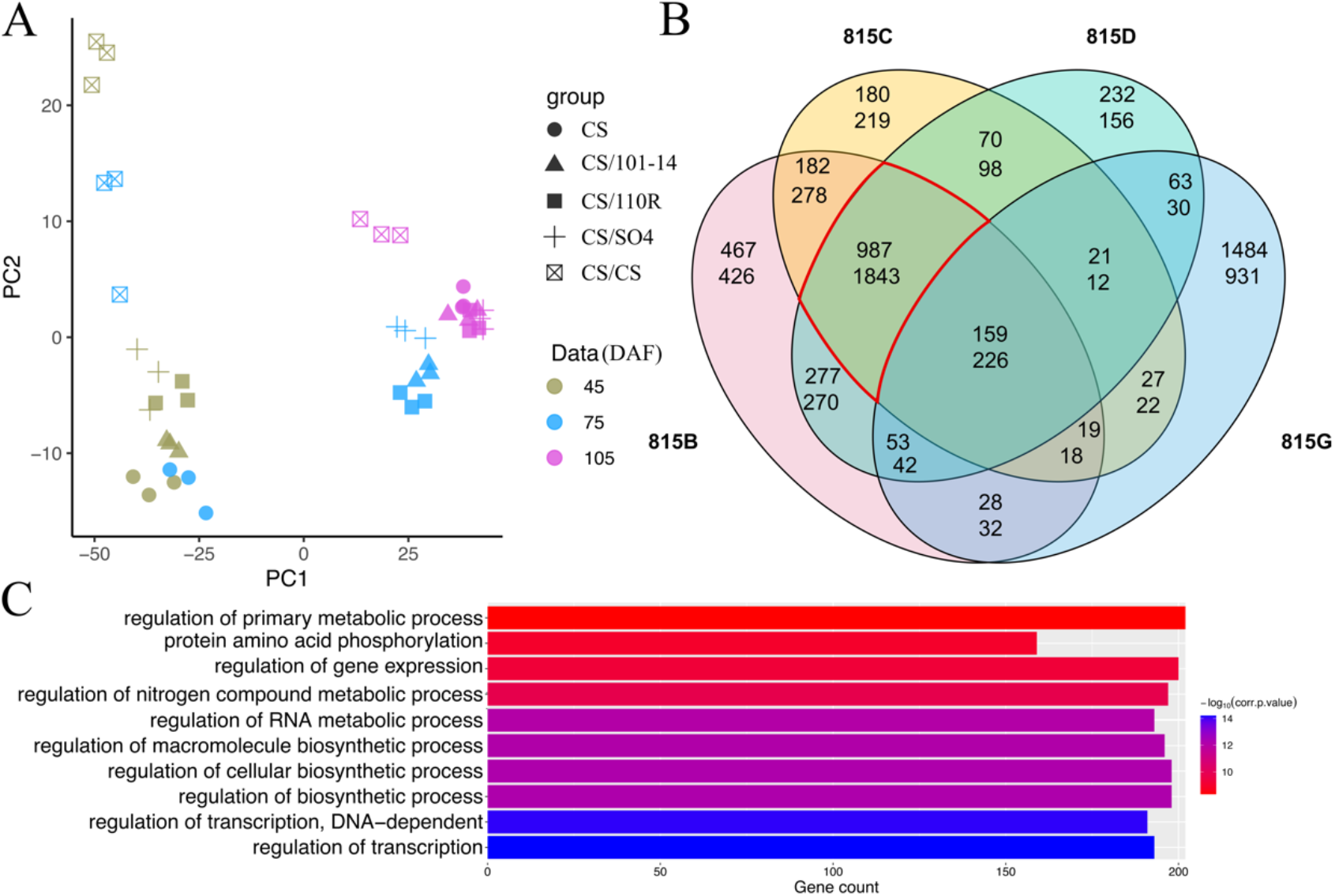
Variability of transcriptional levels among grapes grafted with different rootstocks. A, PCA results of the transcriptome data. B, the Overlap of the DEGs in 4 rootstocks compared with self-root, upper number and lower number means the number of up-regulated down-regulated genes. C, The first ten GO terms enriched in the DEGs of the common part (highlighted in B) of three rootstocks.

The differentially expressed genes (DEGs) screened with |FPKM|>1 and FDR ≤ 0.05 and resulted in 11972 DEGs identified in different stage compared with self-rooted grafting-free (CS) samples, representing 52.27% of the whole-genome transcripts. The number of DEGs in every group ranged from 550-4539, and at the 75 DAF stage, the DEGs number is bigger than other stages (Figure S1). Therefore, the differences of gene expression observed in PCA were well supported by the DEG analyses (Figure 3A). According to the Venn diagram at 75 DAF, the differential expressed genes was 815 CS/CS specific/815 common in the three heterografting samples, with 1484/987 upregulated genes and 931/1843 downregulated genes, respectively (Figure 3B). Further, we categorized the functions by the singular enrichment analysis of the DEGs overlapped in the three heterografting samples at 75 DAF. The results showed 136 and 5 enriched GO terms derived from the upregulated and downregulated DEGs, respectively. A total of 66 of the 136 upregulated terms belonged BP catalogue, in which “regulation of transcription”, “regulation of biosynthetic process”, “regulation of primary process”, “response to abiotic stimulus”, “response to carbohydrate stimulus” and “response to mechanical stimulus” were the most enriched ones (Figure 3C, Table S3).

### Gene screening using the WGCNA analysis

To classify the co-expression modules and identify hub genes based on transcriptomic and metabolomic data, a weighted correlation network was constructed using 25038 transcripts. In the present study, a thresholding power of 3 was selected, which was the lowest power that properly fits the scale-free topological index, and 17 modules revealed after the merged dynamic analysis (Figure S2A). The modules were sorted and numbered according to the gene number assigned to each module. Most of the genes (16948) fell into the first module while the module 2-5 includes genes more than 500, and genes in the other 12 modules were distributed between 252-51.

The correlation coefficients between the modules and anthocyanin content varied widely from −0.73 to 0.92. Four intriguing modules (module 8, 10, 11 and 13) with GS-value greater than 0.5 in multiple compounds or PCs were screened, indicating genes in these modules have significant correlation with the anthocyanin content. The biological functions of the intrinsic genes in the four modules were further analyzed (Figure 4A). First, due to these four modules were in a same cluster, so we analyzed the function of all genes. The result revealed four term related chitin catabolic process, stilbenoid, diarylheptanoid and gingerol biosynthesis, abscisic acid binding, and phenylalanine ammonia-lyase activity (Figure S2B). Next each module were checked and the two modules we focused on were module 8 and module 11. In module 8, five KEGG terms and 14 GO terms were enriched, in which five terms related to ‘Phenylalanine’”, such as KEGG:00940 (Phenylpropanoid biosynthesis) and GO:0009699 (Phenylpropanoid biosynthesis) showed genes in this module participated in the synthesis and metabolism of compounds related anthocyanin. In addition, KEGG:00945 (Stilbenoid, diarylheptanoid, and gingerol biosynthesis) and GO:0009738 (abscisic acid-activated signaling pathway) indicated this module play other roles in berry development. In module 11, only 3 GO term were enriched including DNA-binding transcription activator activity, production of siRNA involved in RNA interference, and gene silencing by RNA, indicating that these modules were mainly associated with the siRNA activities. Connectivity, MM and GS value of genes in each module was calculated and combed to screen the hub genes (Figure S3). In this study, 82, 22, 57 and 43 hub genes were identified in modules 8, 10, 11, and 13 respectively (Figure 5). A total of 16 Hub-TF genes were detected and classified into 8 TF families by using PlantTFDB. Based on Hub-TF genes and correlation network, we built and visualized the network highly related the anthocyanin synthesis (Figure 5).

**Figure 4.**
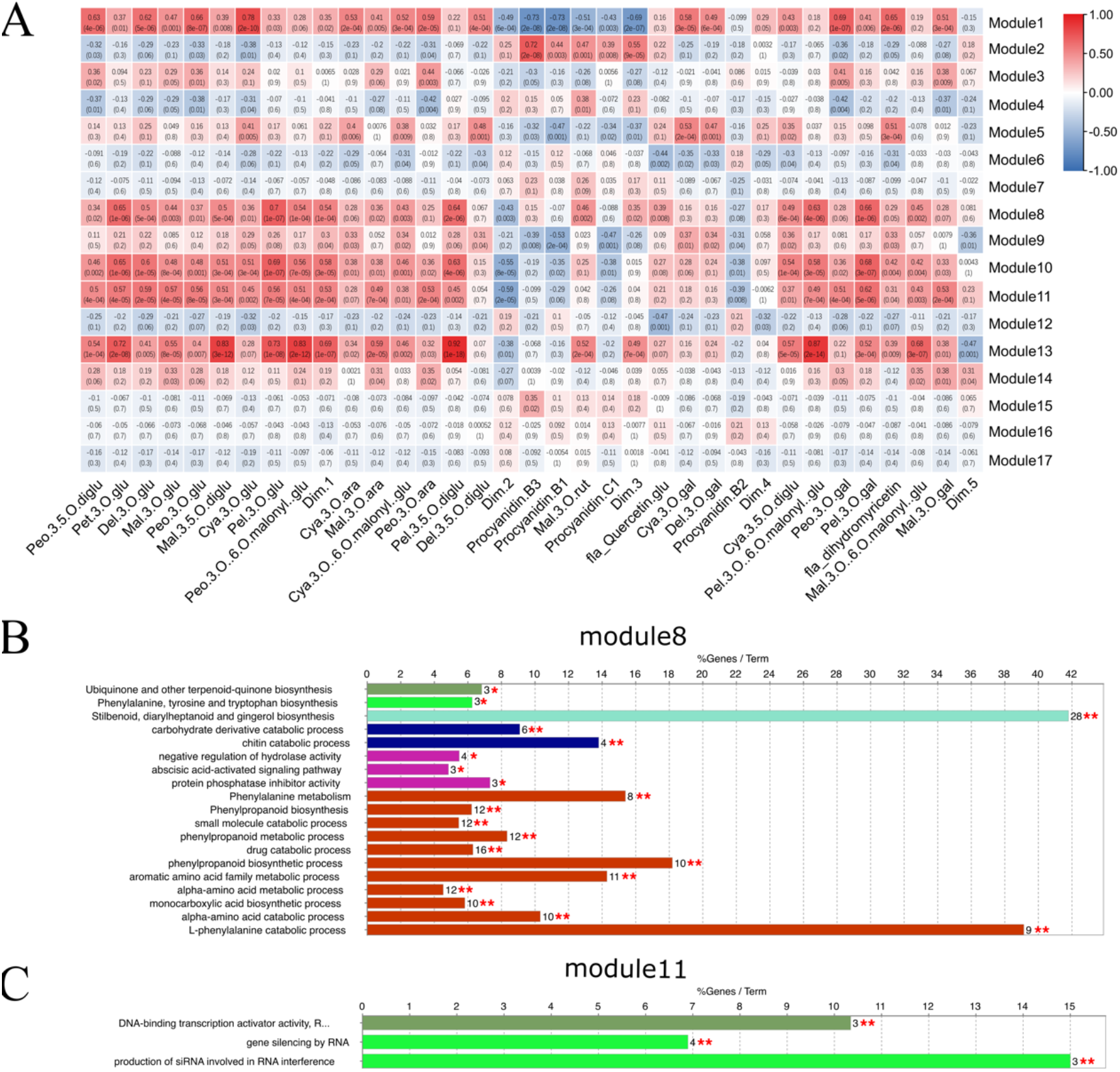
The module-anthocyanin association analysis. A, Heatmap shows the correlation between modules and anthocyanin. Abbreviations and full names correspond to Table S4. The GS-value between a given module and anthocyanin is indicated by the color of the cell and the text inside cells (upper number is the value and lower number is P-value). Red and blue indicated positive and negative correlation, respectively. B and C, The GO-enrichment analysis of Module 8 and 11, respectively. *, P < 0.05; **, P < 0.01; and ***, P < 0.001.

**Figure 5.**
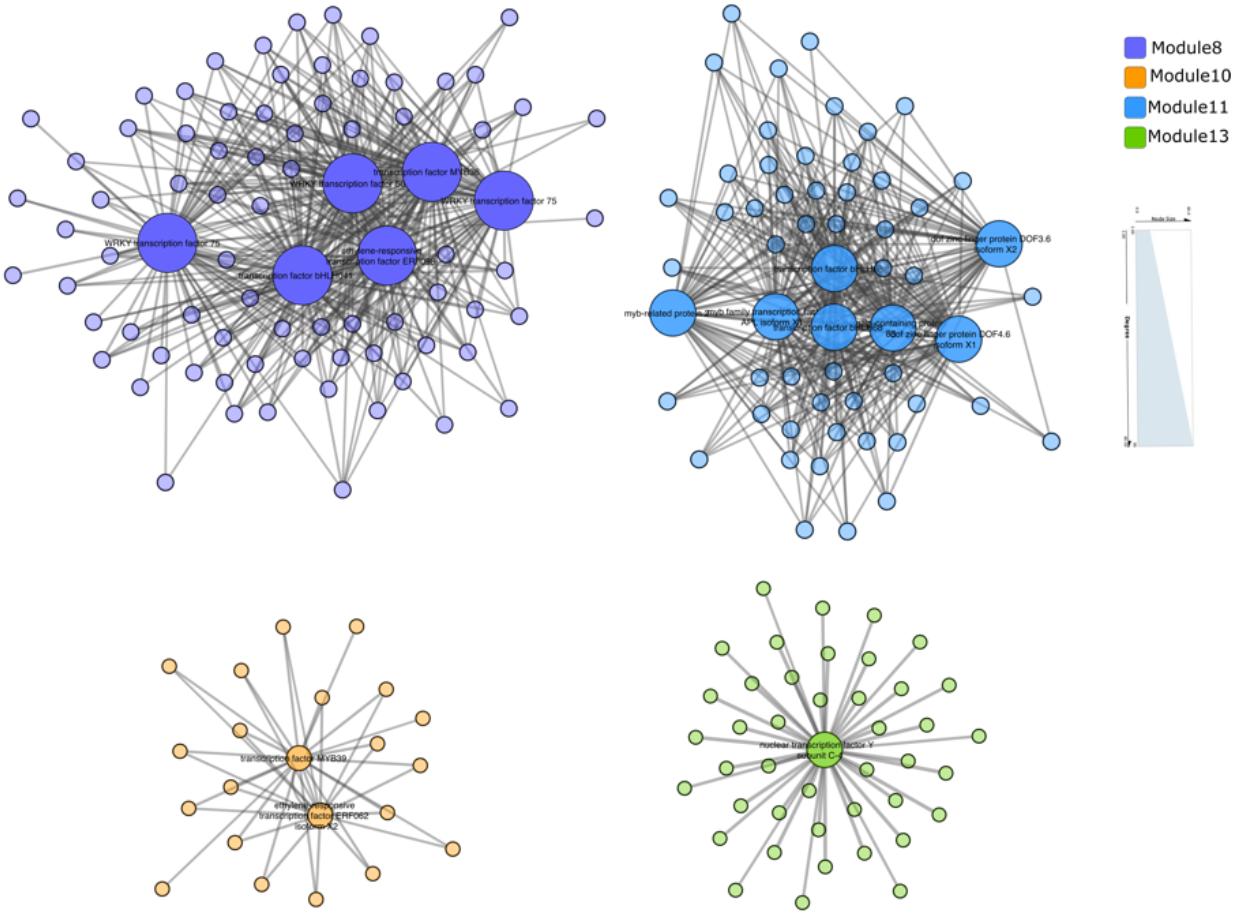
The correlation network in modules highly related to anthocyanin synthesis. The size of node represents the number of genes connected. The transparency of edges means the weight value between two genes.

### The expression pattern and the validation of anthocyanins’ biosynthetic pathway genes

We selected 19 genes in anthocyanin biosynthetic pathway belonged to 11 gene families. We found all the genes expressed differently in all samples (Figure 6, S4). During fruit coloring, most of genes of anthocyanin biosynthetic pathway were upregulated with the highest expression level at 105 DAF. However, each gene family has more than one gene from two genes (DFR) to 13 genes (PAL) with high expression levels. It is indicated that although they might have the similar functions, but only a few genes were functional. Comparing with self-grafting CS/CS and grafting-free CS samples, genes in the rootstock group started upregulated earlier in the former than the later, especially in a *PAL, 4CH, 4CL, CHS, CHI,* and *F3H,* which participate the anthocyanin precursor synthesis in the anthocyanin pathway. And, the same with the trend of phenotype processes and anthocyanin content were observed, the highest gene expression in CS/CS at 105 DAF than the other four groups (Figure 6, S4).

**Figure 6.**
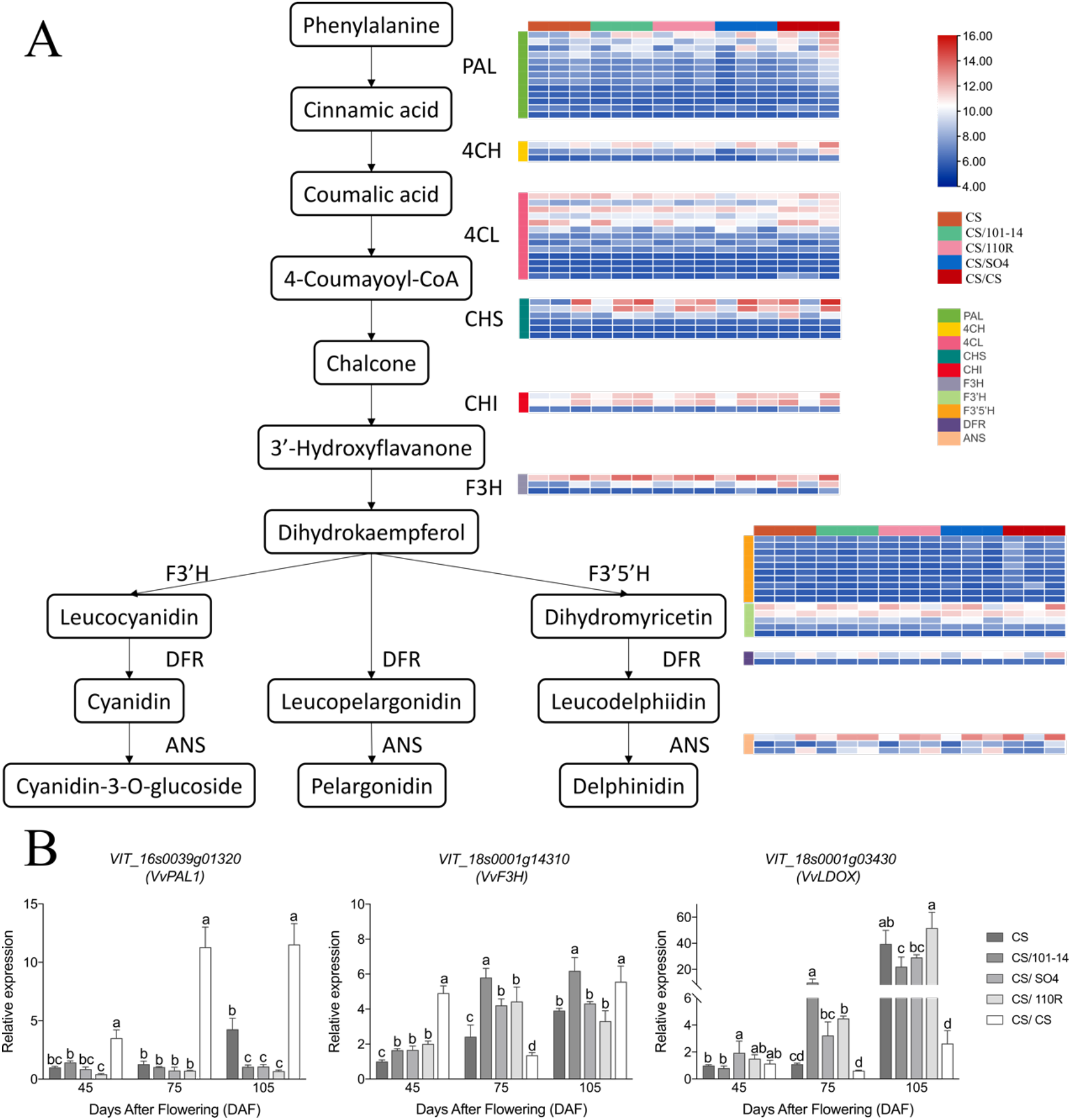
The transcript profiling (A) and RT-PCR(B) of genes in the anthocyanin biosynthetic pathway. Grids with color-scale from blue to white to red represented the RPKM values of DEGs from low to middle to high. PAL, phenylalanine ammonia-lyase; C4H, cinnamic acid 4-hydroxylase; 4CL, 4-coumarate CoA ligase; CHS, chalcone synthase; CHI, chalcone isomerase; F3H, flavanone 3-hydroxylase; F3’H, flavonoid 3’-hydroxylase; F3’5’H, flavonoid 3’,5’-hydroxylase; DFR, dihydroflavonol 4-reductase; ANS, anthocyanidin synthase.

Finally, 19 DEGs including 4 transcription factors, 4 genes of phenylpropanoid biosynthetic pathway, and 12 genes of flavonoid biosynthetic pathway were selected to analyze their expression levels in all samples using RT-qPCR. The results validated the good consistency between RNA-Seq data and RT-qPCR, with a correlation coefficient was 0.9992 (Figure S5).

## Discussion

The grafting in grapevine production can improve the fitness and phenotypes of the scion plant, such berry quality, berry coloring, environmental adaptation, and disease resistance. According to our field observation, the grafting of CS scion with the 101-14MG rootstock had a positive influence on fruit coloring. However, the molecular mechanism at the micro-level is unknown. We combined the berry color phenotypes, metabolomic and transcriptomic data at three stages of berry development of CS grafted to four rootstocks (three heterografting: CS/101-14MG, CS/SO4, CS/110R, and one self-grafting CS/CS) with an own-rooted grafting-free Crimson Seedless (CS) as a control. The results indicated that the heterografting had up-regulated the genes expression that involved in the anthocyanin biosynthesis pathway and promoted an earlier reddening of the berries in CS/101-14MG, CS/SO4, and CS/110R. The TF factors are the hubs in regulation of the early reddening. The self-grafting plants (CS/CS) also showed an earlier reddening, more anthocyanin content and upregulated of genes in the anthocyanin synthesis pathway than the grafting-free plants (CS), suggesting the grafting practice alone might have positive effects on berry reddening in grapevine.

The pigments responsible for the purplish-red color peels of the CS cultivar included Cyanidin 3-O-glucoside, Delphinidin 3-O-glucoside, Malvidin 3-O-glucoside, Peonidin 3-O-glucoside and Petunidin 3-O-glucoside (Figure 2). In the samples of group CS/101-14MG, the content of anthocyanins significantly increased from 75 DAF, while at 105 DAF, the accumulation of anthocyanins in groups CS/SO4 and CS/CS was dramatically higher than that in group CS (Figure 1). The results showed that rootstock grafting could improve the content of anthocyanins in grape berries and promote coloration. The grafting material of rootstock 101-14MG could promote the accumulation of anthocyanins in grape berries in advance.

Previous studies addressed the ‘MYB-bHLH-WDR’ regulatory complex coordinately activated multiple genes of anthocyanin^40,41^. In bright colored fruits, the genes encoding key enzymes downstream of anthocyanin biosynthesis pathway are often highly expressed, such as *DFR, ANS,* and *UFGT*^42^. The MBW complex consisted of MYB transcription factor, basic helix-loop-helix (bHLH), and WD40 proteins was demonstrated to regulate the expression of anthocyanin genes^43^. In *Arabidopsis thaliana,* some MYB transcription factors such as TT2, MYB75, MYB113, and MYB114, some bHLH transcription factors such as TT8, GL3, and EGL3, and a WD40 repeat protein TTG1 can regulate the expression levels of several downstream genes, such as DFR, ANS, and UFGT and affect the anthocyanin biosynthesis^43^.

In this study, the anthocyanins biosynthetic related genes from the upstream (phenylalanine ammonia-lyase, cinnamic acid 4-hydroxylase, 4-coumarate CoA ligase, chalcone synthase, flavanone 3-hydroxylase, flavonoid 3’ -hydroxylase, flavonoid 3’,5 ‘-hydroxylase, flavonoid 3’ -hydroxylase, flavonoid 3’,5 ‘-hydroxylase, and dihydroflavonol 4-reductase) to the downstream (anthocyanidin 3-O-glucosyltransferase and anthocyanidin synthase) were almost upregulated with the accumulating of anthocyanins and berry reddening. However, all these genes were also highly expressed in CS/CS samples, the results suggested that self-grafting rootstocks might have an earlier response to fruit color-related metabolism. The differentially expressed MYBs, such as transcription factor *MYB44* and transcription factor *MYB4* were hubs in PPI interacting network analysis. We predict that MYBs are the key regulators involved in anthocyanin pathways in the interactions between grapevine and rootstocks.

In apple, *CHS* is positively regulated by the expression of *MYB4* and *MYB5*^44^ The *FcMYB1* in strawberry switches the accumulations of anthocyanins and flavonoids on and off^45^. However, the deletion of MYB *cis*-elements in *CHS* promotor can cause white crabapple morphs^46^. In our PPI interacting network analysis, the *trans*-cinnamate 4-monooxygenase-like was directly interaction with transcription factors MYB86 and MYB4; the leucoanthocyanidin reductase 1 was directly interaction with MYBPA1 protein; the flavonoid 3’5’ hydroxylase, anthocyanidin 3-O-glucosyltransferase 2, and MYC anthocyanin regulatory protein were directly interaction with transcription factor MYB90; the flavonoid 3’ hydroxylase was directly interaction with MYB-related protein 308 and MYB-related protein 305.

The DELLA proteins positively regulate the biosynthesis of anthocyanin in *Arabidopsis.* The DELLA proteins can directly interact with and sequester the AtMYBL2 and AtJAZ repressors, resulting in higher MBW complex activities^47^. A considerable number of anthocyanin repressors have been consistently identified. In *Arabidopsis* seedlings, miR858 inhibits the expression of anthocyanin repressor *AtMYBL2,* thus regulating the anthocyanin biosynthesis positively^48^. In tomatoes, inversely, miR858 inhibits the expression of *SlMYB7-like* to regulates anthocyanin biosynthesis negatively. Blocking miR858 function via ectopic expression of a small tandem target mimic of miR858 enhanced anthocyanin accumulation in tomato seedlings^49^. high auxin concentration inhibits anthocyanin biosynthesis^50,51^. A study of red-fleshed apple calli^52^ demonstrated that Auxin Response Factor 13 (MdARF13) inhibited the biosynthesis of anthocyanin. It was achieved both by the direct binding of MdARF13 to the promoter of the ABP gene *MdDFR* to repress its expression and by the physical interaction of MdARF13 with the subgroup 6 R2R3-MYB activator MdMYB10 to destabilize the MBW complex. In PPI interacting network, transcription repressor MYB4-like transcription factor was directly interaction with transcription factor bHLH87, transcription factor bHLH106 and DEAD-box ATP-dependent RNA helicase 42.

DEAD-box ATP-dependent RNA helicase 42 was situated hub of PPI interacting network and directly related to all MYB genes including all MYB transcription factors, MYB-related proteins, *MYBPA2* and transcription factor GAMYB. The DEAD-box RNA helicases participate in ribonucleoprotein complexes rearrangement and RNA structure modification, thereby participating in all aspects of RNA metabolism. The DEAD-box RNA Helicase42 (OsRH42) is necessary to support effective splicing of pre-mRNA during mRNA maturation at low temperatures^53^. The importance of DEADbox ATP-dependent RNA helicase 42 in anthocyanin metabolism was to be expected. The module 11 significantly correlated with anthocyanin content and enriched for siRNA activities and siRNA had played important roles in the regulation networks between the scions and the stocks^3^.

## Conclusions

In summary, the combined phenotypes, transcriptome, and metabolome comprehensive analyses provided large-scale information on gene-metabolite regulation networks related to anthocyanin synthesis. Our results provide global transcriptional changes in grape peel color regulation under different grafting conditions for improving the production and breeding of grapevine.

## Author Contributions

Conceptualization, HZ, ZL, and FZ; Data curation, HZ, ZL, FZ, XZ, XS, WL, HX, NW. Formal analysis, HZ, WL, HX, and NW. Funding acquisition, HZ and MP. Methodology, HZ, ZL, and FZ. Project administration, HZ. Resources, FZ, ZL XZ, and XW; Supervision, MP, XW and YZ. Validation, HZ ZL, and FZ. Writing, review and editing, HZ, ZL, FZ, and YZ.

## Data availability

The RNA-Seq dataset in this study have been deposited in the NCBI under the project number xxxx.

## Acknowledgments

This research was financed by the National Natural Science Foundation of China (32160682), the National Natural Science Foundation of China (31960575), the basic scientific research funding project of Tianshan Youth-Excellence Youth Project (No. 2020Q028) funds project, Supported by China Agriculture Research System of MOF and MARA, Forestry reform and Development from the central government (Xin [2021] TG04).

