## Supplemental files for "Metabolomic and transcriptomic analyses reveal the effects of grafting on anthocyanin synthesis in grapevine"

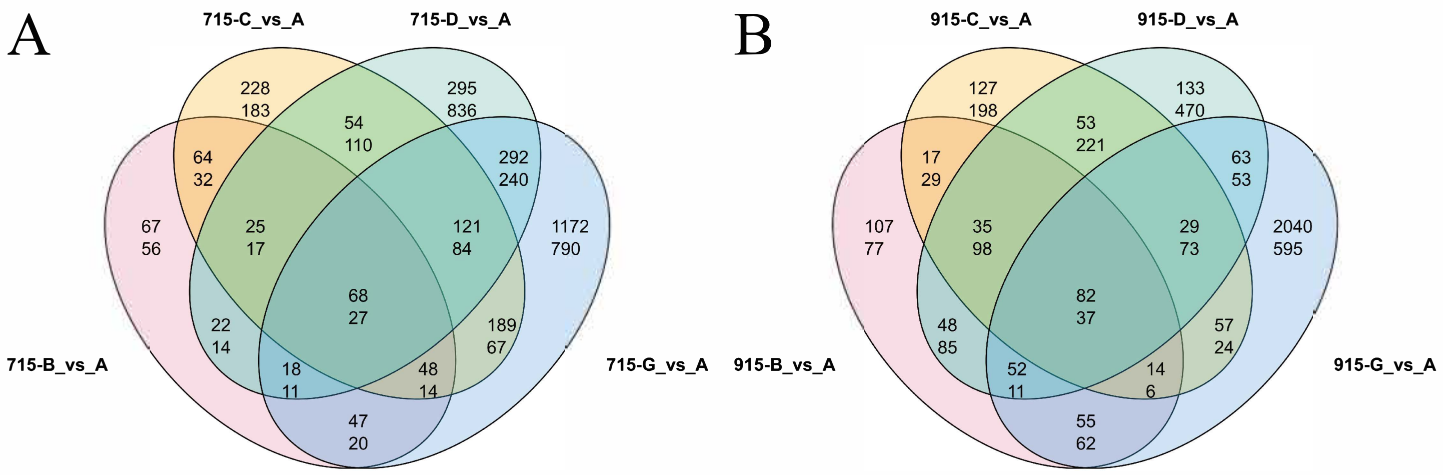


FigureS1 Veen plots showing the overlap of DEGs in 2 stages: 45 DAF(A) and 105DAF (B).
Overlap of the DEGs in 4 rootstocks compared with self-root. Upper number is the gene number of up-regulated and the lower number means down-regulated.

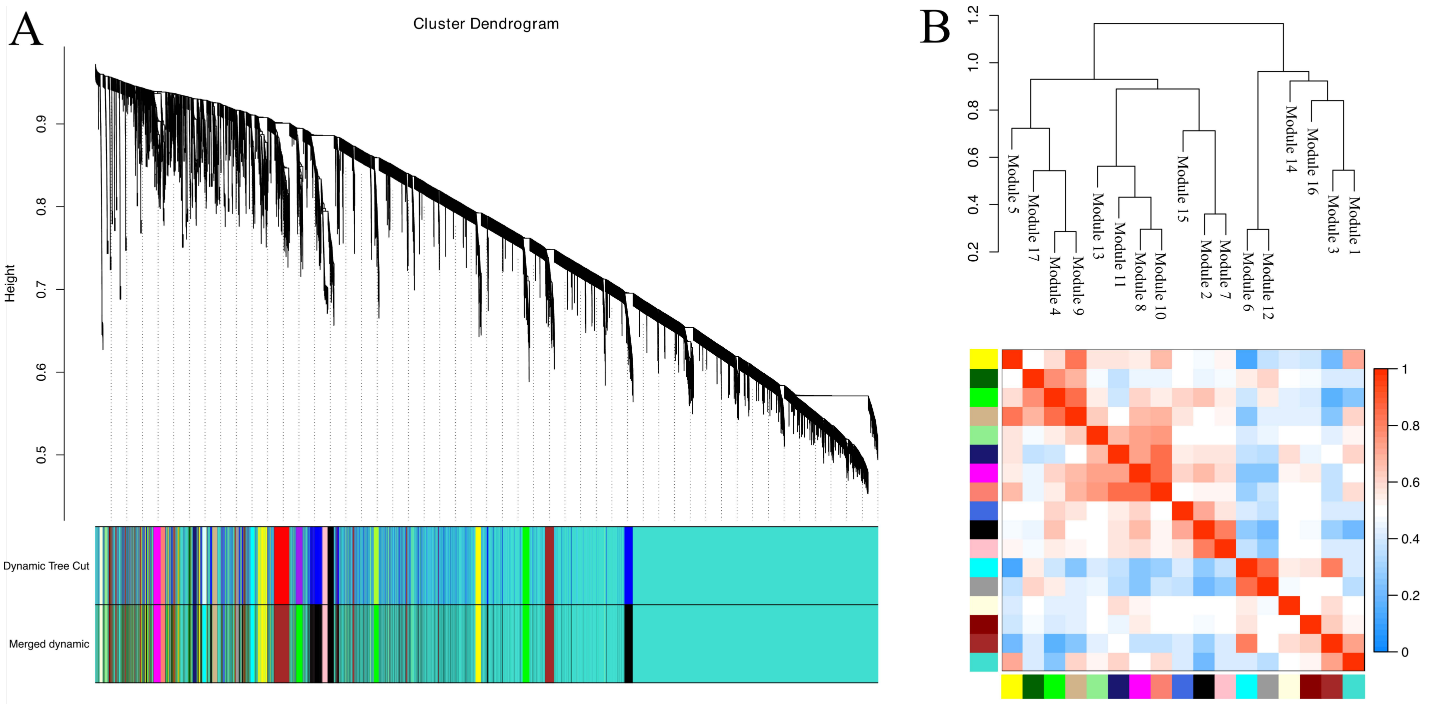


FigureS2 Weighted-gene co-expression network

A Hierarchical cluster dendrogram constructed by WGCNA, on which each leaf represents a gene. 17 merged modules (based on a threshold of 0.20) identified by weighted-gene co-expression network.

B. Module cluster dendrogram and module adjacency heatmap. Cluster dendrogram of module eigengenes. Branches of the dendrogram group together eigengenes that are positively correlated.

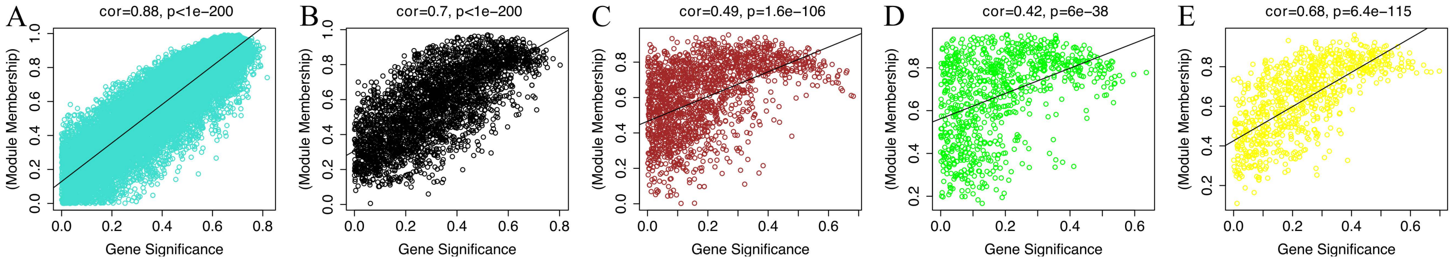


FigureS3 Relationship between module membership (MM) and gene significance (GS)
Scatterplots show the relationship between GS and MM in first 5 modules(A-H: Module 1-5). Illustrating that gene highly significantly associated with a trait are often also the most important (central) elements of modules associated with the trait.

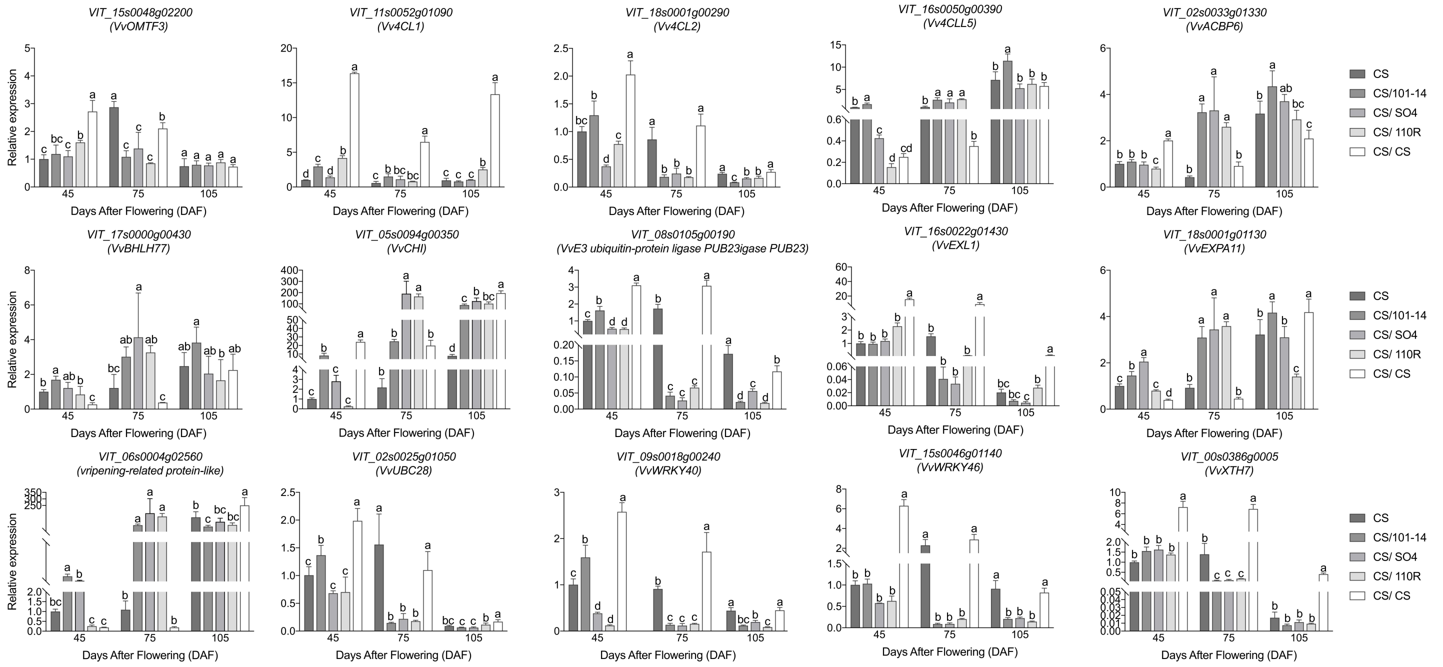


FigureS4 Expression patterns of 15 DEGs in three stages of grape with self-root and grafted on 4 rootstocks. Lowercase letters indicate significance at the 0.01 levels.

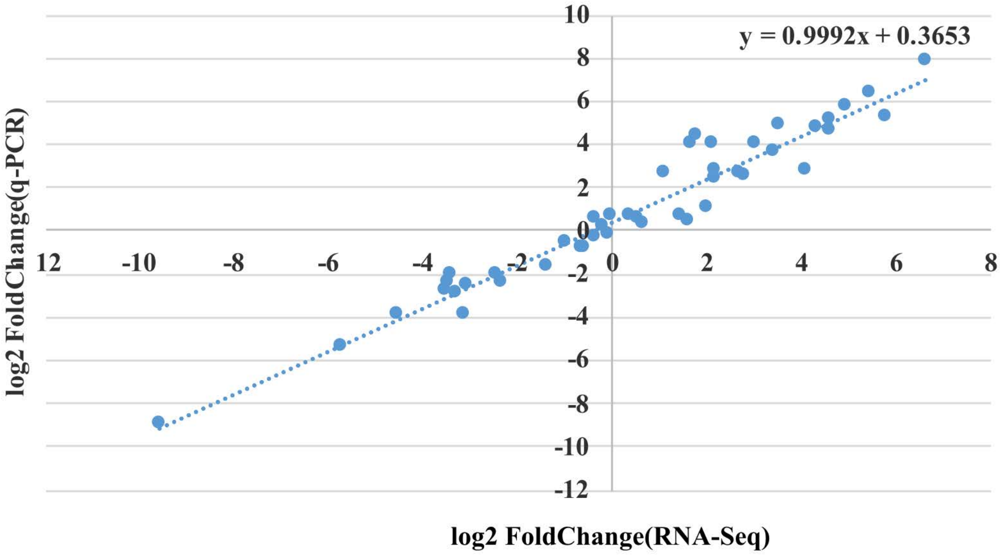


FigureS5 The correlation coefficient diagram of RNA-Seq data and RT-qPCR.

TableS1 List of primers used for RT-qPCR.

| Gene name | Gene ID | Forward primer(3'-5') | Reverse primer(3'-5') |
|---|---|---|---|
| Gapdh | | TTCTCGTTGAGGGCTATTCC | CCACAGACTTCATCGGTGACA |
| CHI | VIT_05s0094g00350 | GGAGTCCTTTTGGCAGCCTT | CACCGGAAGAACCACCACTT |
| ripening-related protein-like | VIT_06s0004g02560 | TTCAGAGCGCATCGTAGCAT | CATGCTCTTTGTCACAGCCG |
| ACBP6 | VIT_02s0033g01330 | CCGTCCGGGGATATTCAACC | TCCTCCAGCAACTGTTTTACCT |
| EXPA11 | VIT_18s0001g01130 | ATGATGGGGCTTCATGTGGG | ACCACCCTCCATTGTTGCTT |
| BHLH77 | VIT_17s0000g00430 | AGGGAACACCACCACTTGTT | AGCCTATCCACCACTGACGA |
| LDOX | VIT_18s0001g03430 | CCCTCTTTGTCCATGTCGGT | ATTGCCTTATGGGGAGGTGC |
| OMTF3 | VIT_15s0048g02200 | GGGGCGGAACGATGTATTGA | CGTGTTTCACGTCCACCTTG |
| F3H | VIT_18s0001g14310 | GAAGCTGATGGGTCTTGCCT | GAGTCCGAGCGTGAGATCAG |
| 4CLL5 | VIT_16s0050g00390 | TCCACGTTCACGGCTTACTC | ATTTGGTGGATCGTGGGGAC |
| EXL1 | VIT_16s0022g01430 | CGGTGACAGCCAAAACGAAG | GCATAGTTTGGCGGAGTTGC |
| UBC28 | VIT_02s0025g01050 | AGTCCATATGCTGGTGGAGTG | ATCGAGAGCAACACCTTGGA |
| E3 ubiquitin-protein ligase PUB23 | VIT_08s0105g00190 | AAAGATCCGGTGACGGTGTC | GGAGAGTGTGGTTGGGAGTC |
| XTH7 | VIT_00s0386g00050 | TGGTGGGAGGGAACTGCATA | TTAGATGCCTGCGACACACT |
| WRKY40 | VIT_09s0018g00240 | CCAGCCTTGTGGTGAAGGAT | TCTTGACAGGGCAGCTAGGA |
| WRKY46 | VIT_15s0046g01140 | CTCCATCATCATCCCGCGAA | TGCCTGCTTGGTCCTTGAAA |
| CHS | VIT_16s0100g00950 | TGTCAAGTGCATGTGTGTTGT | AAGCCTGGTCCAAAACCGAA |
| 4CL1 | VIT_11s0052g01090 | GAGATCGAGAAGCAGGCGAA | TGATTTTCACGGTGGGGAGG |
| 4CLL2 | VIT_18s0001g00290 | AGCAGACCTTCATCTGCACC | GGCTCGGTACTTGGAGATGG |
| PAL1 | VIT_16s0039g01320 | ATCCGCTGAACTGGGGAATG | ACCTGCGATATGGTAAGCGT |

Supplementary Table S2. Raw data, clean data, mapping rate statistics.

| Sample | Reads No. | Q20 (%) | Q30 (%) | Clean Reads No. | Clean Reads % | Total_Mapped | Uniquely_Mapped |
|---|---|---|---|---|---|---|---|
| A715_1 | 45863786 | 97.23 | 92.68 | 41549692 | 90.59 | 38336895 (92.27%) | 37292617 (97.28%) |
| A715_2 | 47693444 | 96.94 | 92.07 | 43129422 | 90.43 | 39526677 (91.65%) | 38475385 (97.34%) |
| A715_3 | 44019828 | 97.36 | 92.95 | 39498884 | 89.72 | 36336197 (91.99%) | 35355849 (97.30%) |
| B715_1 | 46283648 | 97.36 | 92.94 | 42207540 | 91.19 | 37387781 (88.58%) | 36359198 (97.25%) |
| B715_2 | 44357984 | 96.92 | 91.98 | 39893286 | 89.93 | 33388551 (83.69%) | 32497687 (97.33%) |
| B715_3 | 44088268 | 97.24 | 92.67 | 39893592 | 90.48 | 33732096 (84.56%) | 32828546 (97.32%) |
| G715_1 | 45931010 | 96.6 | 91.3 | 41608136 | 90.58 | 37993822 (91.31%) | 37015262 (97.42%) |
| G715_2 | 47027376 | 97.09 | 92.33 | 42749972 | 90.9 | 39285924 (91.90%) | 38250912 (97.37%) |
| G715_3 | 45330362 | 96.85 | 91.82 | 41224562 | 90.94 | 37397851 (90.72%) | 36451641 (97.47%) |
| A815_1 | 46672100 | 96.97 | 92.09 | 42000788 | 89.99 | 38675155 (92.08%) | 37689018 (97.45%) |
| A815_2 | 47653160 | 97.22 | 92.66 | 42444462 | 89.06 | 39177893 (92.30%) | 38143328 (97.36%) |
| A815_3 | 47182260 | 97.45 | 93 | 42591712 | 90.27 | 39471903 (92.68%) | 38434416 (97.37%) |
| B815_1 | 52291752 | 96.79 | 91.69 | 47426754 | 90.69 | 42896148 (90.45%) | 41613525 (97.01%) |
| B815_2 | 46883078 | 96.92 | 92 | 42753778 | 91.19 | 38404311 (89.83%) | 37231297 (96.95%) |
| B815_3 | 45337084 | 97.07 | 92.26 | 41613848 | 91.78 | 37706410 (90.61%) | 36554899 (96.95%) |
| G815_1 | 48653634 | 97.57 | 93.33 | 43922764 | 90.27 | 40654758 (92.56%) | 39607130 (97.42%) |
| G815_2 | 49623402 | 97.33 | 92.77 | 45325058 | 91.33 | 42067104 (92.81%) | 41013340 (97.50%) |
| G815_3 | 45304300 | 97.25 | 92.66 | 40691000 | 89.81 | 37439541 (92.01%) | 36513649 (97.53%) |
| A915_1 | 49786462 | 96.67 | 91.53 | 44929124 | 90.24 | 40394742 (89.91%) | 38951849 (96.43%) |
| A915_2 | 48527366 | 97.03 | 92.23 | 44510200 | 91.72 | 40413763 (90.80%) | 39107314 (96.77%) |
| A915_3 | 46135856 | 97.34 | 92.84 | 42083052 | 91.21 | 38379441 (91.20%) | 37023177 (96.47%) |
| B915_1 | 46613520 | 97.23 | 92.61 | 42828378 | 91.87 | 38978550 (91.01%) | 37762750 (96.88%) |
| B915_2 | 46129072 | 97.06 | 92.3 | 41902250 | 90.83 | 38135277 (91.01%) | 36843800 (96.61%) |

| | | | | | | | |
|---|---|---|---|---|---|---|---|
| B915_3 | 43657302 | 97.17 | 92.47 | 40196426 | 92.07 | 36890726 (91.78%) | 35693041 (96.75%) |
| G915_1 | 49675564 | 97.11 | 92.34 | 45775264 | 92.14 | 42029397 (91.82%) | 40724533 (96.90%) |
| G915_2 | 46287960 | 96.68 | 91.48 | 42187354 | 91.14 | 38025333 (90.13%) | 36760683 (96.67%) |
| G915_3 | 46104048 | 97.38 | 92.93 | 42074676 | 91.26 | 37858710 (89.98%) | 36570549 (96.60%) |

TableS3 GO enrichment analysis of DEGs overlap in BCD at stage 2

**Upregulated**

| GO-ID | p-value | corr p-value | x | n | X | N | Description |
|---|---|---|---|---|---|---|---|
| 4612 | 7.59E-06 | 1.62E-02 | 4 | 5 | 749 | 21148 | phosphoenolpyruvate carboxykinase (ATP) activity |
| 71554 | 2.95E-05 | 2.41E-02 | 30 | 374 | 749 | 21148 | cell wall organization or biogenesis |
| 6732 | 3.40E-05 | 2.41E-02 | 27 | 322 | 749 | 21148 | coenzyme metabolic process |
| 8199 | 9.75E-05 | 4.45E-02 | 4 | 8 | 749 | 21148 | ferric iron binding |
| 16757 | 1.04E-04 | 4.45E-02 | 43 | 662 | 749 | 21148 | transferase activity, transferring glycosyl groups |

**Downregulated**

| GO-ID | p-value | corr p-value | x | n | X | N | Description |
|---|---|---|---|---|---|---|---|
| 3700 | 1.06E-19 | 2.47E-16 | 111 | 642 | 1457 | 21147 | transcription factor activity |
| 30528 | 1.36E-16 | 1.59E-13 | 116 | 753 | 1457 | 21147 | transcription regulator activity |
| 45449 | 6.12E-16 | 4.77E-13 | 193 | 1571 | 1457 | 21147 | regulation of transcription |
| 6355 | 1.42E-15 | 8.31E-13 | 191 | 1562 | 1457 | 21147 | regulation of transcription, DNA-dependent |
| 31326 | 1.50E-13 | 7.03E-11 | 198 | 1718 | 1457 | 21147 | regulation of cellular biosynthetic process |
| 9889 | 1.88E-13 | 7.33E-11 | 198 | 1722 | 1457 | 21147 | regulation of biosynthetic process |
| 10556 | 2.75E-13 | 9.19E-11 | 196 | 1706 | 1457 | 21147 | regulation of macromolecule biosynthetic process |
| 51252 | 3.47E-13 | 1.02E-10 | 193 | 1676 | 1457 | 21147 | regulation of RNA metabolic process |
| 3677 | 1.37E-12 | 3.57E-10 | 193 | 1701 | 1457 | 21147 | DNA binding |
| 19219 | 5.01E-11 | 1.17E-08 | 197 | 1818 | 1457 | 21147 | regulation of nucleobase, nucleoside, nucleotide and nucleic acid metabolic process |
| 51171 | 5.79E-11 | 1.23E-08 | 197 | 1821 | 1457 | 21147 | regulation of nitrogen compound metabolic process |
| 10468 | 2.00E-10 | 3.90E-08 | 200 | 1883 | 1457 | 21147 | regulation of gene expression |
| 6468 | 2.28E-10 | 4.10E-08 | 159 | 1407 | 1457 | 21147 | protein amino acid phosphorylation |

| | | | | | | | |
|---|---|---|---|---|---|---|---|
| 4672 | 2.46E-10 | 4.10E-08 | 158 | 1397 | 1457 | 21147 | protein kinase activity |
| 43565 | 1.10E-09 | 1.71E-07 | 80 | 573 | 1457 | 21147 | sequence-specific DNA binding |
| 80090 | 1.80E-09 | 2.64E-07 | 202 | 1957 | 1457 | 21147 | regulation of primary metabolic process |
| 43687 | 3.54E-09 | 4.87E-07 | 232 | 2340 | 1457 | 21147 | post-translational protein modification |
| 5886 | 1.66E-08 | 2.16E-06 | 207 | 2073 | 1457 | 21147 | plasma membrane |
| 16773 | 2.08E-08 | 2.56E-06 | 163 | 1547 | 1457 | 21147 | phosphotransferase activity, alcohol group as acceptor |
| 16301 | 4.38E-08 | 5.06E-06 | 168 | 1624 | 1457 | 21147 | kinase activity |
| 46777 | 4.69E-08 | 5.06E-06 | 30 | 145 | 1457 | 21147 | protein amino acid autophosphorylation |
| 60255 | 4.76E-08 | 5.06E-06 | 204 | 2064 | 1457 | 21147 | regulation of macromolecule metabolic process |
| 31323 | 7.33E-08 | 7.46E-06 | 210 | 2150 | 1457 | 21147 | regulation of cellular metabolic process |
| 4674 | 3.62E-07 | 3.53E-05 | 100 | 875 | 1457 | 21147 | protein serine/threonine kinase activity |
| 31224 | 6.48E-07 | 6.06E-05 | 481 | 5800 | 1457 | 21147 | intrinsic to membrane |
| 6464 | 9.56E-07 | 8.60E-05 | 240 | 2607 | 1457 | 21147 | protein modification process |
| 31407 | 1.12E-06 | 9.67E-05 | 13 | 39 | 1457 | 21147 | oxylipin metabolic process |
| 16310 | 1.32E-06 | 1.10E-04 | 169 | 1724 | 1457 | 21147 | phosphorylation |
| 9768 | 1.67E-06 | 1.35E-04 | 9 | 19 | 1457 | 21147 | photosynthesis, light harvesting in photosystem I |
| 9522 | 2.13E-06 | 1.66E-04 | 13 | 41 | 1457 | 21147 | photosystem I |
| 16772 | 2.44E-06 | 1.84E-04 | 173 | 1791 | 1457 | 21147 | transferase activity, transferring phosphorus-containing groups |
| 19222 | 2.51E-06 | 1.84E-04 | 216 | 2333 | 1457 | 21147 | regulation of metabolic process |
| 16021 | 3.73E-06 | 2.65E-04 | 469 | 5719 | 1457 | 21147 | integral to membrane |
| 44425 | 5.86E-06 | 4.03E-04 | 508 | 6281 | 1457 | 21147 | membrane part |
| 9694 | 1.15E-05 | 7.69E-04 | 9 | 23 | 1457 | 21147 | jasmonic acid metabolic process |
| 16020 | 1.32E-05 | 8.58E-04 | 591 | 7487 | 1457 | 21147 | membrane |
| 16740 | 1.50E-05 | 9.52E-04 | 311 | 3643 | 1457 | 21147 | transferase activity |
| 50982 | 2.24E-05 | 1.38E-03 | 4 | 4 | 1457 | 21147 | detection of mechanical stimulus |
| 4871 | 2.63E-05 | 1.54E-03 | 33 | 223 | 1457 | 21147 | signal transducer activity |

| | | | | | | | |
|---|---|---|---|---|---|---|---|
| 60089 | 2.63E-05 | 1.54E-03 | 33 | 223 | 1457 | 21147 | molecular transducer activity |
| 6796 | 3.04E-05 | 1.73E-03 | 186 | 2036 | 1457 | 21147 | phosphate metabolic process |
| 6793 | 3.13E-05 | 1.74E-03 | 186 | 2037 | 1457 | 21147 | phosphorus metabolic process |
| 44459 | 3.45E-05 | 1.88E-03 | 59 | 499 | 1457 | 21147 | plasma membrane part |
| 7166 | 4.56E-05 | 2.43E-03 | 35 | 249 | 1457 | 21147 | cell surface receptor linked signaling pathway |
| 4872 | 4.91E-05 | 2.54E-03 | 29 | 191 | 1457 | 21147 | receptor activity |
| 18298 | 4.99E-05 | 2.54E-03 | 10 | 33 | 1457 | 21147 | protein-chromophore linkage |
| 9765 | 5.13E-05 | 2.55E-03 | 9 | 27 | 1457 | 21147 | photosynthesis, light harvesting |
| 50794 | 5.31E-05 | 2.59E-03 | 248 | 2863 | 1457 | 21147 | regulation of cellular process |
| 4888 | 5.73E-05 | 2.73E-03 | 28 | 183 | 1457 | 21147 | transmembrane receptor activity |
| 4675 | 8.12E-05 | 3.65E-03 | 27 | 177 | 1457 | 21147 | transmembrane receptor protein serine/threonine kinase activity |
| 7167 | 8.12E-05 | 3.65E-03 | 27 | 177 | 1457 | 21147 | enzyme linked receptor protein signaling pathway |
| 7178 | 8.12E-05 | 3.65E-03 | 27 | 177 | 1457 | 21147 | transmembrane receptor protein serine/threonine kinase signaling pathway |
| 9521 | 8.68E-05 | 3.81E-03 | 18 | 96 | 1457 | 21147 | photosystem |
| 44212 | 8.86E-05 | 3.81E-03 | 38 | 288 | 1457 | 21147 | DNA regulatory region binding |
| 19199 | 8.96E-05 | 3.81E-03 | 27 | 178 | 1457 | 21147 | transmembrane receptor protein kinase activity |
| 31408 | 9.58E-05 | 3.97E-03 | 8 | 23 | 1457 | 21147 | oxylipin biosynthetic process |
| 9611 | 9.67E-05 | 3.97E-03 | 9 | 29 | 1457 | 21147 | response to wounding |
| 4702 | 1.02E-04 | 4.04E-03 | 28 | 189 | 1457 | 21147 | receptor signaling protein serine/threonine kinase activity |
| 5057 | 1.02E-04 | 4.04E-03 | 28 | 189 | 1457 | 21147 | receptor signaling protein activity |
| 10158 | 1.06E-04 | 4.14E-03 | 4 | 5 | 1457 | 21147 | abaxial cell fate specification |
| 31225 | 1.08E-04 | 4.15E-03 | 20 | 115 | 1457 | 21147 | anchored to membrane |
| 50789 | 1.10E-04 | 4.15E-03 | 266 | 3134 | 1457 | 21147 | regulation of biological process |
| 9734 | 1.12E-04 | 4.15E-03 | 14 | 65 | 1457 | 21147 | auxin mediated signaling pathway |
| 16168 | 1.30E-04 | 4.74E-03 | 9 | 30 | 1457 | 21147 | chlorophyll binding |
| 71365 | 1.33E-04 | 4.78E-03 | 14 | 66 | 1457 | 21147 | cellular response to auxin stimulus |

| | | | | | | | |
|---|---|---|---|---|---|---|---|
| 31347 | 1.40E-04 | 4.96E-03 | 12 | 51 | 1457 | 21147 | regulation of defense response |
| 4842 | 1.73E-04 | 6.01E-03 | 24 | 156 | 1457 | 21147 | ubiquitin-protein ligase activity |
| 46658 | 1.75E-04 | 6.01E-03 | 19 | 110 | 1457 | 21147 | anchored to plasma membrane |
| 6857 | 1.87E-04 | 6.24E-03 | 8 | 25 | 1457 | 21147 | oligopeptide transport |
| 15833 | 1.87E-04 | 6.24E-03 | 8 | 25 | 1457 | 21147 | peptide transport |
| 16881 | 1.89E-04 | 6.24E-03 | 27 | 186 | 1457 | 21147 | acid-amino acid ligase activity |
| 9523 | 2.56E-04 | 8.32E-03 | 14 | 70 | 1457 | 21147 | photosystem II |
| 10200 | 3.08E-04 | 9.70E-03 | 6 | 15 | 1457 | 21147 | response to chitin |
| 9922 | 3.26E-04 | 9.70E-03 | 3 | 3 | 1457 | 21147 | fatty acid elongase activity |
| 34625 | 3.26E-04 | 9.70E-03 | 3 | 3 | 1457 | 21147 | fatty acid elongation, monounsaturated fatty acid |
| 34626 | 3.26E-04 | 9.70E-03 | 3 | 3 | 1457 | 21147 | fatty acid elongation, polyunsaturated fatty acid |
| 19367 | 3.26E-04 | 9.70E-03 | 3 | 3 | 1457 | 21147 | fatty acid elongation, saturated fatty acid |
| 19368 | 3.26E-04 | 9.70E-03 | 3 | 3 | 1457 | 21147 | fatty acid elongation, unsaturated fatty acid |
| 30154 | 3.27E-04 | 9.70E-03 | 31 | 233 | 1457 | 21147 | cell differentiation |
| 6629 | 3.46E-04 | 1.01E-02 | 85 | 851 | 1457 | 21147 | lipid metabolic process |
| 19787 | 4.07E-04 | 1.18E-02 | 24 | 165 | 1457 | 21147 | small conjugating protein ligase activity |
| 10287 | 4.45E-04 | 1.27E-02 | 8 | 28 | 1457 | 21147 | plastoglobule |
| 9582 | 5.01E-04 | 1.41E-02 | 5 | 11 | 1457 | 21147 | detection of abiotic stimulus |
| 31226 | 5.10E-04 | 1.42E-02 | 31 | 239 | 1457 | 21147 | intrinsic to plasma membrane |
| 9506 | 5.27E-04 | 1.43E-02 | 30 | 229 | 1457 | 21147 | plasmodesma |
| 55044 | 5.27E-04 | 1.43E-02 | 30 | 229 | 1457 | 21147 | symplast |
| 30054 | 5.67E-04 | 1.51E-02 | 30 | 230 | 1457 | 21147 | cell junction |
| 5911 | 5.67E-04 | 1.51E-02 | 30 | 230 | 1457 | 21147 | cell-cell junction |
| 23033 | 6.55E-04 | 1.72E-02 | 82 | 832 | 1457 | 21147 | signaling pathway |
| 9695 | 6.63E-04 | 1.72E-02 | 4 | 7 | 1457 | 21147 | jasmonic acid biosynthetic process |
| 9581 | 8.11E-04 | 2.08E-02 | 5 | 12 | 1457 | 21147 | detection of external stimulus |

| | | | | | | | |
|---|---|---|---|---|---|---|---|
| 16298 | 8.27E-04 | 2.10E-02 | 16 | 96 | 1457 | 21147 | lipase activity |
| 5509 | 1.25E-03 | 3.12E-02 | 32 | 263 | 1457 | 21147 | calcium ion binding |
| 42761 | 1.25E-03 | 3.12E-02 | 4 | 8 | 1457 | 21147 | very long-chain fatty acid biosynthetic process |
| 16879 | 1.63E-03 | 4.02E-02 | 28 | 224 | 1457 | 21147 | ligase activity, forming carbon-nitrogen bonds |
| 250 | 1.77E-03 | 4.32E-02 | 6 | 20 | 1457 | 21147 | lanosterol synthase activity |
| 38 | 2.13E-03 | 4.76E-02 | 4 | 9 | 1457 | 21147 | very long-chain fatty acid metabolic process |
| 8381 | 2.13E-03 | 4.76E-02 | 4 | 9 | 1457 | 21147 | mechanically-gated ion channel activity |
| 9612 | 2.13E-03 | 4.76E-02 | 4 | 9 | 1457 | 21147 | response to mechanical stimulus |
| 42391 | 2.13E-03 | 4.76E-02 | 4 | 9 | 1457 | 21147 | regulation of membrane potential |
| 30322 | 2.13E-03 | 4.76E-02 | 4 | 9 | 1457 | 21147 | stabilization of membrane potential |
| 22833 | 2.13E-03 | 4.76E-02 | 4 | 9 | 1457 | 21147 | mechanically gated channel activity |
| 22840 | 2.13E-03 | 4.76E-02 | 4 | 9 | 1457 | 21147 | leak channel activity |
| 22841 | 2.13E-03 | 4.76E-02 | 4 | 9 | 1457 | 21147 | potassium ion leak channel activity |
| 22842 | 2.13E-03 | 4.76E-02 | 4 | 9 | 1457 | 21147 | narrow pore channel activity |
| 4620 | 2.17E-03 | 4.79E-02 | 12 | 68 | 1457 | 21147 | phospholipase activity |
| 9743 | 2.19E-03 | 4.79E-02 | 8 | 35 | 1457 | 21147 | response to carbohydrate stimulus |
| 6633 | 2.29E-03 | 4.96E-02 | 18 | 125 | 1457 | 21147 | fatty acid biosynthetic process |

Table S4 Abbreviations and full names correspond in Figure 4

| Abbreviations | Full name |
|---|---|
| Cya-3-O-glu | Cyanidin 3-O-glucoside |
| Del-3-O-gal | Delphinidin 3-O-galactoside |
| Del-3-O-glu | Delphinidin 3-O-glucoside |
| Peo-3-O-glu | Peonidin 3-O-glucoside |
| Procyanidin B1 | Procyanidin B1 |
| Procyanidin B2 | Procyanidin B2 |
| Procyanidin B3 | Procyanidin B3 |
| Procyanidin C1 | Procyanidin C1 |
| fla_Quercetin-glu | Quercetin 3-O-glucoside |
| Pet-3-O-glu | Petunidin 3-O-glucoside |
| Del-3,5-O-diglu | Delphinidin 3,5-O-diglucoside(Delphin) |
| Cya-3-O-gal | Cyanidin 3-O-galactoside |
| Mal-3-O-glu | Malvidin 3-O-glucoside |
| Peo-3,5-O-diglu | Peonidin 3,5-O-diglucoside |
| Mal-3-O-rut | Malvidin 3-O-rutinoside |
| Mal-3,5-O-diglu | Malvidin 3,5-diglucoside |
| Peo-3-O-gal | Peonidin 3-O-galactoside |
| Mal-3-O-gal | Malvidin 3-O-galactoside |
| Pel-3-O-glu | Pelargonidin 3-O-glucoside |
| Cya-3-O-(6-O-malonyl)-glu | Cyanidin 3-O-(6-O-malonyl-beta-D-glucoside) |
| Cya-3-O-ara | Cyanidin 3-O-arabinoside |
| Pel-3-O-gal | Pelargonidin 3-O-galactoside |
| Peo-3-O-(6-O-malonyl)-glu | Peonidin 3-O-(6-O-malonyl-beta-D-glucoside) |
| Peo-3-O-ara | Peonidin 3-O-arabinoside |
| Mal-3-O-ara | Malvidin 3-O-arabinoside |
| Pel-3,5-O-diglu | Pelnidin 3,5-O-diglucoside |
| Cya-3,5-O-diglu | Cyanidin 3,5-O-diglucoside |
| fla_dihydromyricetin | Dihydromyricetin |
| Pel-3-O-(6-O-malonyl)-glu | Pelargonidin 3-O-(6-O-malonyl-beta-D-glucoside) |
